# Development of a Targeted Choroidal Injury Model for the Study of Retinal Degenerations and Therapeutic Cell Replacement

**DOI:** 10.1101/2025.07.29.667466

**Authors:** Narendra Pandala, Lorena De Melo Haefeli, Mallory Lang, Edwin M. Stone, Robert F. Mullins, Budd A. Tucker, Ian C. Han

## Abstract

**Purpose:** Choroidal loss is an important pathophysiological step in many retinal diseases, but few reliable translational models of choroidal injury exist. Here, we report a new targeted choroidal injury model using bioconjugated saporins and compare it models of systemic sodium iodate administration.

**Methods:** Wild-type Sprague-Dawley rats were given suprachoroidal injections of anti-CD38 or anti-CD105 antibodies conjugated to saporin immunotoxin (10 µl at 0.05 µg/µL) to induce selective choroidal endothelial cell injury. These animals were compared to wild-type rats given sodium iodate (75 mg/kg) via tail vein injections, with a dose escalation study (25, 50, and 75 mg/kg) in immunocompromised (Sprague-Dawley Rag2/Il2g double-knockout) rats. Animals were examined at 1-, 2-, and 3-weeks post-treatment, and the degree of choroidal injury compared using fundus photography, optical coherence tomography, and immunohistochemistry.

**Results:** Suprachoroidal administration of anti-CD38 or anti-CD105 saporins resulted in severe choroidal vascular injury localized to the injection site, without damage to adjacent choroidal vasculature, progressive injury over time, or development of choroidal neovascularization. By contrast, sodium iodate treated animals had rapid, diffuse choroidal loss which progressed throughout the study time points, with fatal systemic side effects at the highest (75 mg/kg) dose.

**Conclusions:** Suprachoroidal injection of anti-CD38 and anti-CD105 saporins results in targeted, localized, non-progressive choroidal injury in rats. These models offer alternatives to systemic sodium iodate administration, which causes diffuse, progressive choroidal injury.

**Translational Relevance:** Immunotoxin-based models of targeted choroidal injury may be useful for understanding pathways of retinal degeneration and facilitating development of therapies for diseases involving choroidal cell loss.

## Introduction

The choroid is a sophisticated vascular structure that provides vital nutrients to the photoreceptors by diffusion as well as removal of waste from the outer retina, thus enabling proper retinal function for vision ^1^. Loss of the choroid is a crucial pathophysiologic step in a wide range of retinal diseases, including common conditions such as age-related macular degeneration (AMD), and rarer inherited retinal diseases such as choroideremia ^2^. The mechanisms of choroidal loss in AMD and other retinal degenerations are complex, involving pathways such as lipofuscin accumulation, oxidative injury, inflammation, and complement activation ^3–7^. The deposition of complement in the choriocapillaris is believed to be a major contributing factor to choroidal degeneration, including in geographic atrophy (GA) in AMD ^4^.

Recently, complement inhibitors delivered by intravitreal injection have emerged as a potential therapy for slowing the progression of GA in AMD ^8–10^. Although these treatments may reduce the rate of GA growth in some patients, their primary goal is preventing further retinal and choroidal injury rather than replacing cells that have already been damaged. Thus, there remains a critical need to pursue other potential therapeutic avenues, including cell replacement strategies designed to restore viable cells to areas of tissue loss, thus offering potential for preserved or improved visual function. To this end, our group has described methods of generating induced pluripotent stem cell (iPSC)-derived choroidal endothelial cells using human skin derived fibroblasts ^11–13^ toward the development of choroidal cell replacement therapy.

One of the current limitations in developing such a choroidal cell replacement approach is the lack of a reliable choroidal injury model to study potential transplantation strategies. Existing models of choroidal destruction rely on ablative injury to the choroid including with laser photocoagulation ^14,15^. While such models are useful for studying treatment of resultant choroidal neovascularization, laser injury results in focal, acute, and severe destruction of multiple structures (e.g., outer retina, Bruch’s membrane, inner choroid). As such, this acute model of retinal and choroidal injury does not parallel the gradual pathophysiologic steps seen in AMD and other retinal degenerations. Moreover, the incited outer retinal scarring from laser injury is less amenable to cell replacement strategies for studying retinal degenerations.

In this study, we describe the development of new models of choroidal endothelial cell injury, including with local suprachoroidal injection of immunotoxins compared to systemic administration of sodium iodate. We demonstrate the feasibility of both widespread and regional choroidal injury in these approaches *in vivo* using a rat model. Among other potential applications, these models will facilitate investigations in choroidal endothelial cell replacement, including with iPSC-derived choroidal endothelial cell transplants.

## Materials and Methods

### Ethics Statement

All rat experiments were conducted with the approval of the Animal Care and Use Committee at the University of Iowa (Animal welfare assurance, A3021-01) and were consistent with the Association for Research in Vision and Ophthalmology (ARVO) Statement for use of animals in ophthalmic and vision research.

### Dosage and time points

Saporin is a plant-derived toxin that induces cell death by acting as a ribosome inactivating protein. To specifically target endothelial cells with saporin, antibodies directed against the cluster of differentiation domains (CD) CD38 and CD105 (proliferating endothelial cells) were used. Anti-CD38 and anti-CD105 antibodies conjugated to saporin were obtained from Advanced Targeting Systems (Carlsbad, CA) and were diluted in sterile PBS at a concentration of 0.05 µg/µl. To induce selective choroidal cell injury, 10 µl of bioconjugate solution was delivered via suprachoroidal injection as described below. An equal number of male and female 6-week-old Sprague Dawley (SD) rats (Charles River Laboratories Wilmington, MA; n=4, per time point) were tested for each immunotoxin injected.

For comparison to the suprachoroidal injection model, sodium iodate (NaIO_3_; Sigma Aldrich, St. Louis, MO) was diluted in 0.9% saline solution, to the requisite concentration, sterile filtered, and administered via tail vein injections, as previously described ^16^. Wild-type Sprague Dawley Rats, 6-8 weeks old with equal number of males and females were used for the NaIO_3_ time course study (n=4, per time point).

To determine the optimal dosing for NaIO_3_ and compare the degree of choroidal injury in an immunocompromised versus wild-type rat model, we performed a similar study as above using 6 to 8-week-old SRG rats (Sprague-Dawley Rag2/Il2rg double-knockout rats; Charles River Laboratories Wilmington, MA; n=4, per dose) with equal number of males and females.

For each method of choroidal injury, rats were evaluated at 1-, 2-, and 3-weeks post-administration with clinical examination, imaging, and immunohistochemistry as further described below.

### Suprachoroidal Injection

Suprachoroidal injections were performed by a fellowship-trained vitreoretinal surgeon (I.C.H.) using techniques as previously described ^17,18^. In brief, the animals were placed under anesthesia using 3-5% inhalant isofluorane gas (Piramal Healthcare, Bethlehem, PA) and their eyes dilated using 1% tropicamide (Alcon Laboratories, Fort Worth, TX). Under direct visualization using an operating microscope, a limited temporal conjunctival peritomy was created using 0.12 forceps and Vannas scissors (Bausch and Lomb/Storz Ophthalmics, Rochester, NY). A 30-gauge needle was used to create a slit in the sclera to access the suprachoroidal space, and a 33-gauge blunt-tipped Hamilton syringe (Hamilton Company, Reno, NV) was used to slowly inject 10 µL of drug (i.e., Anti-CD38-saporin or Anti-CD105-saporin) into the suprachoroidal space. Eyes were examined immediately post-injection to confirm successful creation of a suprachoroidal bleb and absence of complications (e.g., perforation into the subretinal space or vitreous cavity).

### Fundus Examination and Imaging

Animals were examined at various time points post-injection, as outlined above. Prior to sacrifice, eyes were examined under an operating microscope to record clinical findings, including any complications such as inflammation, cataract formation, or retinal detachment. Fundus photography was obtained using a rodent-specific fluorescent fundus camera (Micron IV, Phoenix Laboratories, Pleasanton, CA) to identify the injection site and evaluate for chorioretinal atrophy. Optical coherence tomography (OCT) was also performed (OCT; Phoenix Micron Image-Guided OCT2, Phoenix Laboratories, Pleasanton, CA) to evaluate retinal and choroidal tissue loss.

### Vascular labeling, tissue processing and imaging

The choroidal vasculature of the animals was stained using a modification of a previously described protocol ^19^. Initially, the rats were deeply anesthetized by intraperitoneal injection of ketamine (91 mg/kg) and xylazine (9.1 mg/kg) and placed on a heating pad during the staining procedure. Dylight 488-Tomato Lectin (*Lycopersicon esculentum* agglutinin; Vector Laboratories, Burlingame, CA) was diluted in 0.9% saline solution to a final concentration of 0.1 mg/ml, and 0.5 ml of this solution was injected via intracardiac injection. 5 mins after the cardiac injection, the animals were euthanized using euthasol (150 mg/kg). The eyes were then enucleated and fixed in 4% paraformaldehyde (PFA) in phosphate-buffered saline (PBS) at 4°C for 5 hours. The anterior segment was dissected and removed, and the posterior segment containing the retina and the choroid-sclera complex was flowered using radial cuts into quarters for flat mounting. The retina was then removed, and the choroid-sclera complex was whole mounted onto a glass slide with mounting media and a cover glass. The whole mounts were then examined using a fluorescence microscope (BZ-X810, Keyence, Osaka, Japan). Multiple z-stacks spanning the entire whole mount in x, y and z direction were taken at 20X magnification with 5 µm separation between the images in each z-stack. These raw images were stitched using Keyence image analyzer and Fiji ^20^.

## Results

### Anti-CD105-saporin induced choroidal loss

Saporin is a plant-derived toxin that induces cell death by acting as a ribosome inactivating protein following endocytosis of cell surface ligands ^21^. Here we used immunotoxins (saporin-conjugated antibodies) that bind to endothelial cells (CD105 and CD38). We previously published preliminary data exploring this route of focal choroidal injury ^22^ and showed that anti-CD105 tagged saporin can be used to induce choroidal cell loss in Sprague Dawley (SD) rats. In our prior study, we tested multiple doses via suprachoroidal injections and found out that at higher doses (0.25 µg/µl and 0.5 µg/µl of immunotoxin solution; 10 µl per eye) the toxin diffuses into the overlying layers of the retina causing damage, but at the lower dose of 0.5 µg/eye the damage is localized in the suprachoroidal space, causing choroidal endothelial cell death. Accordingly, in this study, we used that lower dose (0.5 µg/eye) and analyzed the vascular damage at different time points (**Figure 1, Supplementary Figure 1**).

**Figure 1.**
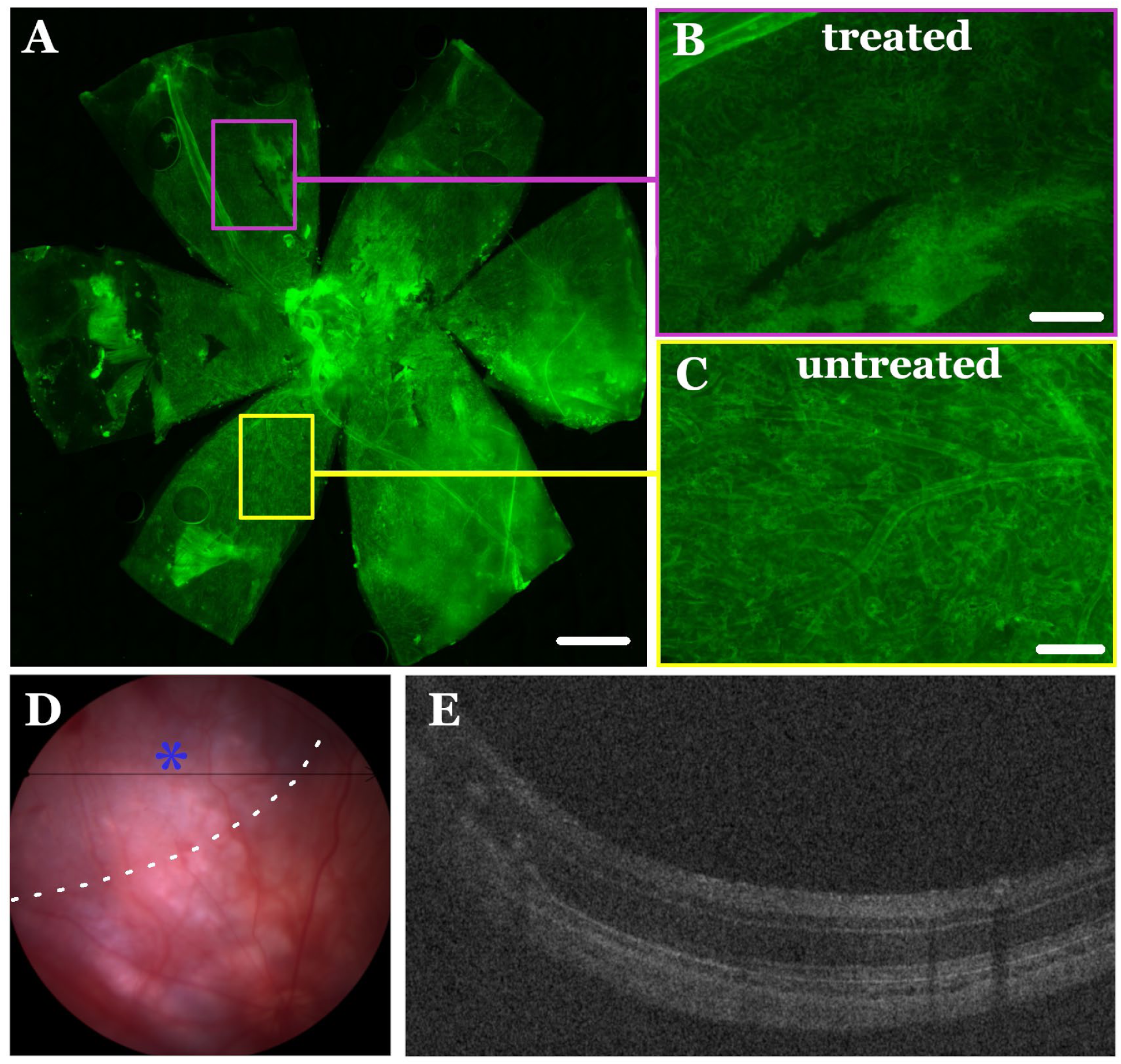
Suprachoroidal injection of anti-CD105-saporin causes sectoral choroidal and outer retinal loss. Suprachoroidal injection of anti-CD105-saporin (0.5 µg/eye) causes sectoral choroidal and outer retinal loss compared to eyes injected with control at day 7. High magnification inserts (B,C) from flat mounts (A; Scale bar is 1000 µm) demonstrate severe choroidal injury with near complete loss of choroidal vessels in treated areas (B; Scale bar is 100 µm) compared to preserved choroidal vasculature in untreated areas (C; Scale bar is 100 µm). Corresponding fundus photographs, with the site of injection denoted by asterisk and outlined in white dashed lines (D), and OCT line scans (E) show sectoral regions of choroidal and outer retinal damage.

Anti-CD105-saporin solution was injected into the suprachoroidal space of wild type SD rats, and the animals were analyzed using fundus photography (**Figure 1 D**) and optical coherence tomography (OCT) (**Figure 1 E**). The choroidal vasculature in the animals was labeled as previously described using fluorophore conjugated tomato lectin via intracardiac injection [15] and high-resolution images of the whole mounted choroid-scleral complex were collected (**Figure 1 A-C**). As shown, suprachoroidal immunotoxin-induced injury was localized to the area of injection, causing loss of the vasculature in this area (**Figure 1 B**), whereas other regions of the choroid remained intact (**Figure 1 C**).

Over the three-week post-injection period, there was persistent damage to the choroidal vasculature without progressive loss and spread of injury to untreated regions (**Supplementary Figure 1**). Clinical examination and imaging revealed no appreciable intraocular inflammation, cataract development, nor retinal injury at any time point studied. No eyes developed choroidal neovascularization (CNV) after treatment. All animals survived the study through the length of the experiment, and no animals were noted to have signs of systemic toxicity nor behavioral changes throughout the study.

### Anti-CD38-saporin induced targeted choroidal loss

For comparison to the anti-CD105-saporin model described above, we assessed the efficacy of anti-CD38-saporin via suprachoroidal injections. CD38 is highly expressed in endothelial cells ^23,24^ whereas CD105 is expressed only in proliferating endothelial cells ^25,26^. As such, we hypothesized that an anti-CD38 immunotoxin moiety may be more effective in targeting all endothelial cells rather than just the proliferating endothelial cells. To test this, SD rats were injected with 0.5 µg of CD38-saporin per eye via suprachoroidal injections, and the effect was assessed at different time points (**Figure 2, Supplementary Figure 2**).

**Figure 2.**
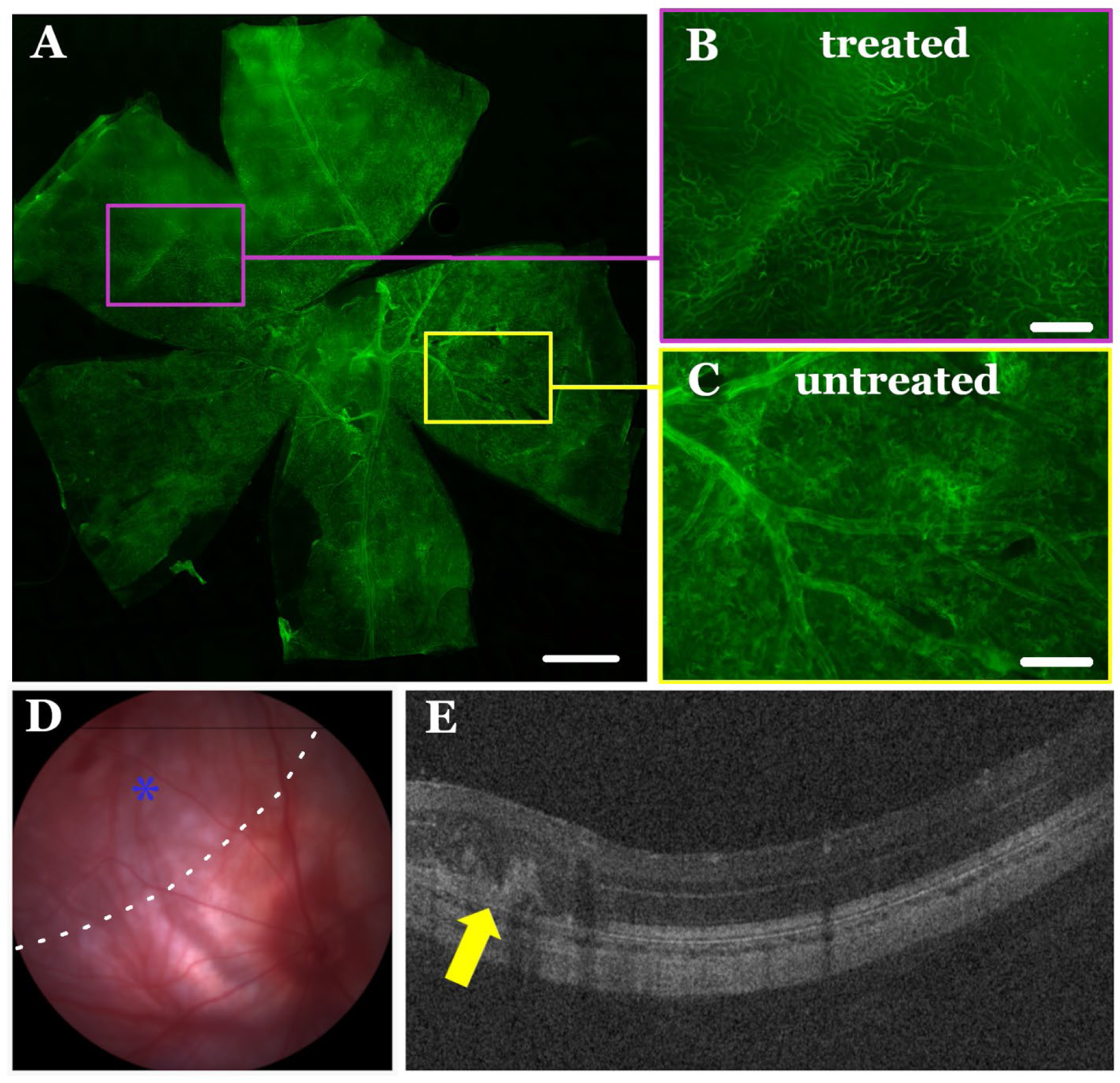
Suprachoroidal injection of anti-CD38-saporin causes sectoral choroidal and outer retinal loss. Anti-CD38-saporin suprachoroidal injection leads to sectoral loss of choroidal and outer retinal tissue (day-7). High magnification images (A; Scale bar is 1000 µm) from choroid scleral flat mounts reveal localized choroidal damage in the treated regions (B; Scale bar is 100 µm), while the choroidal vasculature remains intact in untreated areas (C; Scale bar is 100 µm). Corresponding fundus images, with the site of injection denoted by asterisk and outlined by white dashed lines (D), and OCT scans (E) display distinct sectors with choroidal and outer retinal damage.

Similar to the anti-CD105 immunotoxin injected eyes, suprachoroidal injection of anti-CD38-saporin induced local choroidal injury in the injected region only (**Figure 2 B**). The choroid and retina remote from the injection site remained intact (**Figure 2 C**). As observed in anti-CD105 injected animals, eyes treated with suprachoroidal anti-CD38-saporin had persistent choroidal loss in the treated area through the duration of the 21-day post-injection period (**Supplementary Figure 2**), without qualitative differences in terms of the amount or extent of damage in the vasculature or the outer retina in comparing the groups. No eyes developed CNV after treatment. As with the anti-CD105-saporin animals, all rats treated with anti-CD38-saporin survived the study without visible ocular complication or systemic toxicity.

### Sodium iodate induced choroidal loss

Sodium Iodate (NaIO_3_) induced retinal degeneration is a previously described animal model due its rapid and intense RPE and photoreceptor damage ^16,27^. Prior reports of NaIO_3_ induced choroidal degeneration have predominantly been in pigmented mice ^28,29^. Here we used this model to study the damage to the choroidal vasculature in wildtype SD rats and compare it to the immunotoxin models described above. 75mg/kg of Sodium iodate was administered intravenously, and the animals were studied over multiple time points (**Figure 3**). Based on the flat mount images, the choroidal vascular injury was visible by 1-week post-treatment (**Figure 3 B,F**) and progressively increased over the following two-weeks (**Figure 3 A-H**). At 3 weeks, only the larger blood vessels were present in the choroid, with barely any remaining choriocapillaris (**Figure 3 D,H**), representative of the diffuse, rapid, and progressive choroidal destruction in this model. The NaIO_3_ induced damage was uniform throughout the fundus rather than regions of focal or patchy injury. Over the course of 3 week follow up, there was progressive damage of the overlying retina, with loss of the RPE and vertical, hyperreflective spikes into the outer retina on OCT imaging (**Figure 3 N-P**). No intraocular inflammation or cataract formation was observed in any eyes of NaIO_3_ treated animals, and no eyes developed CNV after treatment. However, of the 12 animals (6 males and 6 females) treated with 75 mg/kg of NaIO_3_, one female rat died after 2 days due to renal failure (hematuria) as a systemic complication of treatment.

**Figure 3.**
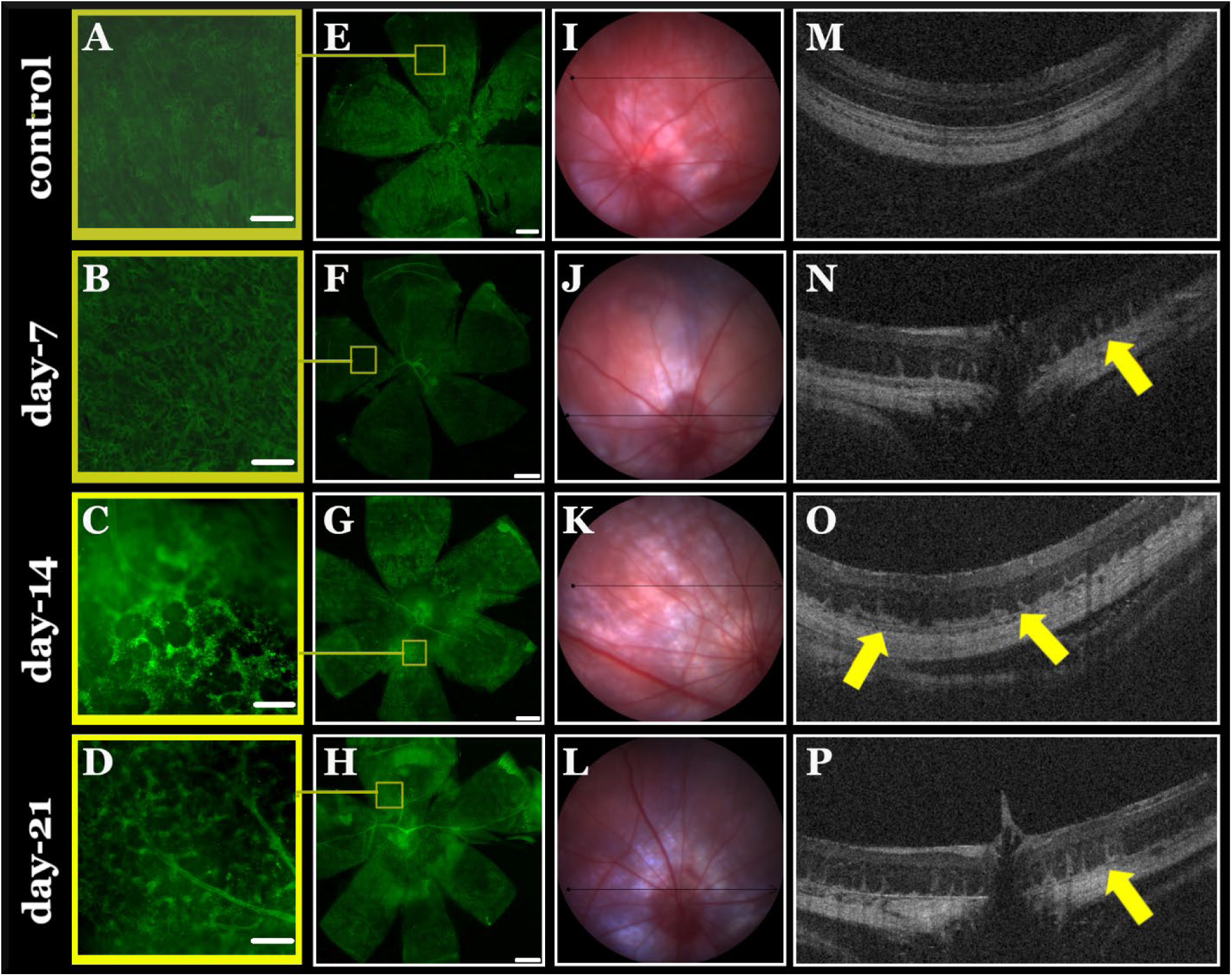
Sodium iodate induces diffuse, progressive choroidal and outer retinal damage. Examples of sodium iodate mediated ablation of choroidal endothelial cells in the Sprague Dawley rat following tail vein injection (75 mg/kg). High magnification insets (A-D, Scale bar is 100 µm) from flat mounts (E-H; Scale bar is 1000 µm) show diffuse choroidal loss at day-7, -14, and -21 post-injection relative to control (top row). Corresponding fundus photographs (I-L) and OCT line scans (M-P) demonstrate diffuse choroidal loss with progressive outer retinal atrophy (yellow arrows).

### Sodium Iodate dosing in SRG animals

One potential application of animal models of choroidal injury is to explore therapeutic cell replacement using human endothelial cell transplantation. To subvert the effects of the xenografts in these transplantation studies, immune compromised animals are needed, but the impact of choroidal injury in these animals is unknown. Previously, we showed the feasibility of hydrogel-supported human induced pluripotent stem cell-derived choroidal endothelial cell transplantation in SRG rats, including those with choroidal damage from CD105 immunotoxin ^22^. Immunocompromised SRG rats lack the lymphoid lineage immune cells (T-, B-, and Natural Killer cells). In this study, we hypothesized that the effect of the choroidal injury in an immunocompromised animal model may be more severe when compared to wild type SD rats. To this end, we tested different doses (25, 50 and 75 mg/kg) of systemic NaIO_3_ and analyzed the retinal and choroidal damage (**Figure 4, Supplementary Figure 3**). By 1-week post-treatment, the choroidal vasculature in SRG rats treated with NaIO_3_ was severely thinned at all 3 doses tested, with a dose-response seen of more severe loss of the choroid and damage to the overlying outer retina at 50 mg/kg and 75 mg/kg. On qualitative comparison to NaIO_3_ treatment in wild-type SD rats, there was more severe choroidal and outer retinal thinning at all doses tested by 1 week post-treatment. Of the 12 animals treated (2M and 2F per dose), one female rat that received the highest (75 mg/kg) dose died after 2 days, with hematuria suggesting renal failure as a systemic complication of treatment.

**Figure 4.**
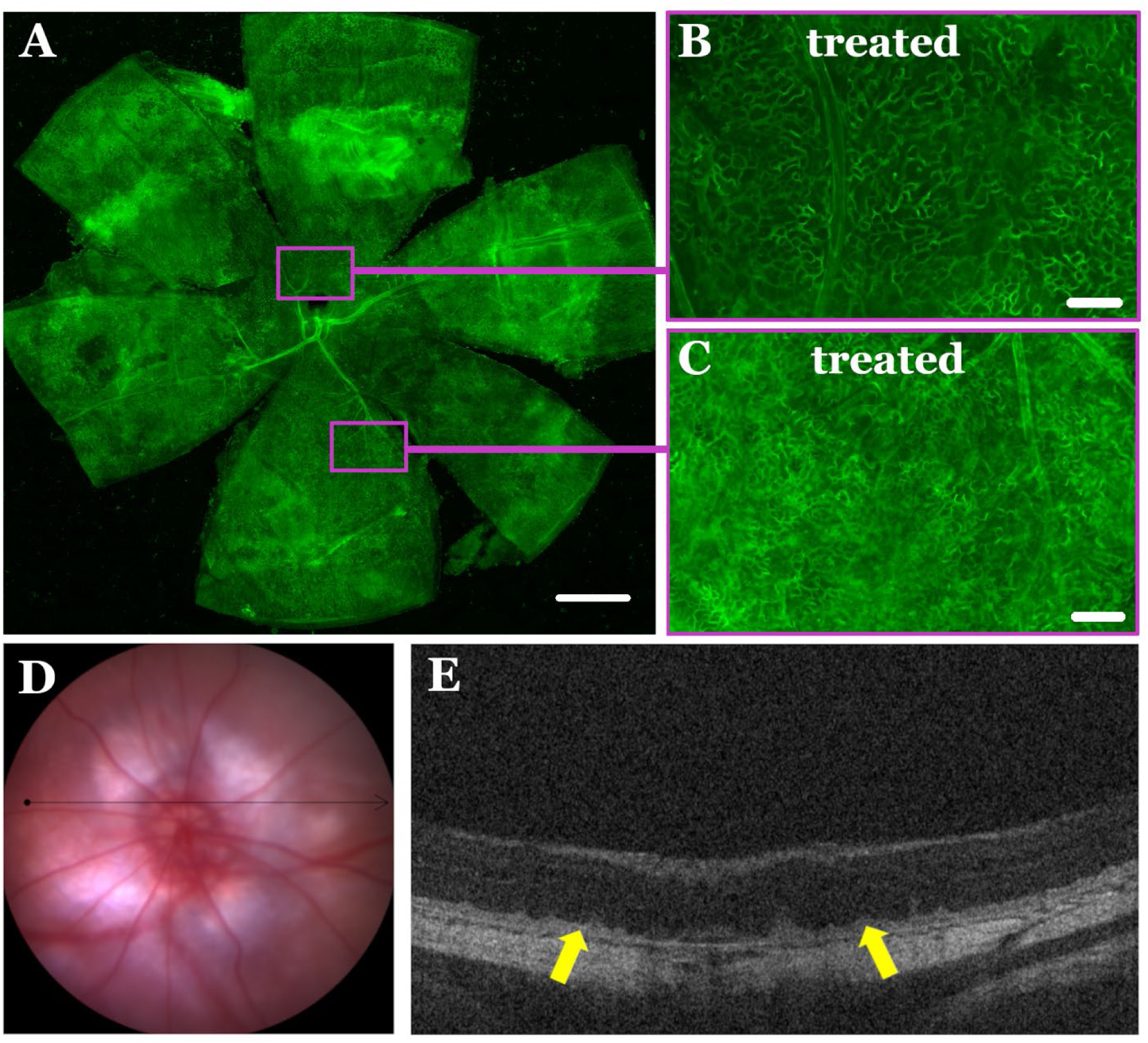
Systemic administration of sodium iodate in immunocompromised SRG rats results in dose-related choroidal and outer retinal damage at 1-week post-treatment. Intravenous sodium iodate resulted in diffuse choroidal and outer retinal loss in a dose-related fashion (75 mg/kg shown here) by 1-week after treatment. High magnification inserts (B, C; Scale bar is 100 µm) from flat mounts (A; Scale bar is 1000 µm) demonstrate choroidal injury. Corresponding fundus photographs (D) and OCT line scans (E) show diffuse choroidal and outer retinal damage (yellow arrows).

## Discussion

In this study, we describe development of a new animal model for targeted choroidal injury using anti-C38-saporin and anti-CD105-saporin delivered via suprachoroidal injection. We show that this approach allows for localized regional injury to the choroid and overlying outer retina with durable effects up to at least 3-weeks post-treatment, without inciting CNV growth. To compare to previously published models, we also evaluated systemic NaIO_3_ administration in both wild-type and immunocompromised SRG rats, demonstrating that there is a dose-related response to diffuse choroidal and outer retinal injury that is more severe in SRG animals than WT, with progressive tissue loss over time, and notable risk of systemic morbidity at higher doses. Because of the reliable, durable, and reproducible choroidal damage (either focal with anti-CD38/105-saporin or more diffuse for NaIO_3_), these models expand the options available for investigations into the pathophysiology of choroidal and retinal degeneration secondary to choroidal endothelial loss, as well as for studies of cell replacement strategies.

A particularly innovative aspect of our study was the use of saporins, ribosome inactivating proteins, to facilitate targeted choroidal injury. Saporins have been used in biological sciences in the past four decades as a versatile molecule to create targeted toxins with applications in Alzheimer’s research, pain studies, and targeted cancer therapies ^30^. Saporin is used in “molecular surgery” for targeted delivery to cells, causing cell death by inhibiting protein synthesis ^30^. Saporin is an extremely stable protein, resistant to denaturation and proteolysis even at temperatures up to 55◦C ^31^. It is also very amenable to coupling with binding proteins and antibodies ^30^. However, there has been limited prior work exploring the utility of saporins in ophthalmic research.

Here we used bioconjugates of saporin to specifically target endothelial cells and create a model of vascular damage by suprachoroidal injections. By conjugating saporin to endothelial cell specific (CD38 and CD105) binding antibodies, we were able to create focal choroidal atrophy in the injected area without any widespread retinal damage. Previously, we also observed some retinal damage induced by the saporin at higher doses, despite the CD105 conjugation. We suspected this was related to the stability of the saporin molecule, which might still be diffusing in the ocular tissue, including outer retina, after choroidal loss ^22^. At the lower doses presented in this paper, we did not see any progressive effect of the saporin, in both the choroid and the adjacent retina, i.e., the damage was contained in the injected area. In addition to a desired local effect of treatment, we did not observe signs of ocular inflammation, CNV formation, or systemic toxicity in the treated animals.

To date, there has been limited work exploring the role of CD38 and CD105 in choroidal endothelial cell survival and function. However, work in other fields has investigated these endothelial cell markers as potential therapeutic targets. For example, daratumumab, a monoclonal antibody that targets CD38, is used to treat multiple myeloma ^32^. Provocatively, some reports have described an association between daratumumab treatment and ocular side effects, including acute angle-closure glaucoma, myopic shift, and choroidal effusions, implying a drug effect on choroidal vasculature in some patients ^33,34^. Post-marketing surveillance and literature review have highlighted the need for immediate ophthalmologic evaluation and intervention upon onset of ocular symptoms ^34^. The exact mechanism for this response is not well understood. However, CD38 ligation has been shown to trigger release of the cytokines IL-1β, IL-6, and IL-10 in resting human monocytes. This occurs through a signaling pathway independent of Fcγ receptors, emphasizing the role of CD38 surface density in monocyte activation ^35^, potentially leading to choroidal effusion due to inflammation in the choroid. CD105 (endoglin) is a cell surface glycoprotein (180 kD) endothelial marker of proliferation ^25,26^. Due to its high expression on the endothelial cells of tumor vasculature, CD105 is a potential target for antibody-based cancer therapies ^25^. CD105 is also expressed highly in the CNV formed in wet AMD, suggesting a potential target for therapeutic interventions in neovascular AMD ^36^, but this remains to be thoroughly explored.

In this study, we compared the novel anti-CD38-saporin and anti-CD105-saporin approaches with a well-described model using systemic NaIO_3_ treatment. NaIO_3_-induced retinal damage was first described in 1941 ^37^, and its effect on the RPE and photoreceptor degeneration has been well studied ^16,38–43^. There are several advantages of this model, including the availability of the chemical and bioavailability through multiple routes of administration over a range of doses ^16,38–41^ from 25 mg/kg to 100 mg/kg ^16,40,41^. Also, the effects of injury can be observed in short span, where photoreceptor swelling is observed within 6 hours of administration and a total loss of retina within 6 months of treatment ^40^. Doses as high as 100 mg/kg have also been used in mice, with RPE cell necrosis followed by photoreceptor apoptosis resulting in a lobular or mosaic pattern of loss, similar to gyrate atrophy of the choroid and retina ^41^. NaIO_3_ has also been administered in the subretinal space, resulting in a significant degeneration of RPE and the photoreceptors, offering a potential model for studying GA ^38,39^.

Despite more published experience with NaIO_3_ models of choroidal and retinal damage, there are also several significant drawbacks to this model for investigators to consider. Specifically, NaIO_3_ has an LD50 of 108 mg/kg IV and 119mg/kg IP ^42^, with significant risk of systemic toxicity and other organ failure particularly at high doses ^16,42^, which we observed in our studies as well. Also, the mechanism of injury is not directly targeted to the choroid. Instead, the oxidative stress induced by NaIO_3_ affects the RPE and photoreceptors first, with subsequent choroidal loss. Interestingly, this mechanism of injury influences the utility of this model for studying CNVs, which are not reliably induced by laser photocoagulation in Long-Evans rats that received treatment with NaIO_3_ ^43^. Also, unlike our model of anti-CD38-saporin or anti-105-saporin via suprachoroidal injection, the choroidal and retinal damage with NaIO_3_ is diffuse throughout the fundus rather than focal and targeted for choroidal injury specifically.

Other rat models of systemic vascular injury exist, including the spontaneously hypertensive (SHR) rat, which is well described particularly for cardiovascular research applications. The SHR rat has been reported to exhibits signs of retinal and choroidal vasculature damage ^44,45^. However, this strain of rats develops hypertension at 5-6 weeks of age and then they start showing the first signs of decreased ocular endothelial surface molecules at 7-8 months of age ^44,46^. Prior publications have used 18-24 month old SHR rats in cardiovascular studies, since they show signs of heart failure and endothelial cell damage ^47^. This might be a good model to study how the endothelial cells and choroid effect the health of retinal layers, but maintaining animal colonies for 18 months is resource intensive.

In summary, we have developed multiple methods of inducing choroidal vascular damage that is readily visible on both clinical imaging and immunohistochemistry. Immunotoxin induced damage in the suprachoroidal space is ideal for studying the focal impact of choroidal cell loss on the adjacent tissue. The local effect of targeted choroidal injury also allows for efficient use of the untreated areas of the same eye as internal controls for comparison (i.e., rather than comparing changes across separate animals with systemic NaIO_3_ model). However, limitations of this model include the need for suprachoroidal injection, which is more technically challenging than systemic intravenous injection with NaIO_3_ and may be more subject to variability across eyes depending on the quality of the injection. Although we noted similar extent and severity of choroidal and retinal injury between anti-CD38-saporin or anti-CD105-saporin treated eyes, we did not perform assessments of retinal or choroidal thickness or degree of vascular loss to quantify differences in the two approaches. The focal choroidal injury in both SAP-related models was reliable and durable across all eyes studied, without CNV formation in any eye; however, we evaluated eyes up until 3 weeks post-injection, and it is possible that the durability of these effects may change over time, or that vasculature remodeling or CNV formation may take place at more distant time points after treatment. Comparatively, the NaIO_3_ model induces diffuse choroidal damage and outer retinal loss with systemic injection via the tail vein, which as indicated above is less technically challenging than suprachoroidal injections. Another advantage of NaIO_3_ is its availability at high purity through commercially available means, whereas bioconjugation of saporins involves more complex chemical synthesis that depends on the antibody or protein to be bound (i.e., greater possibility of batch-to-batch variation, with potential downstream impact on its treatment effect). Future studies may leverage these models of choroidal and outer retinal injury to explore mechanisms of disease related to choroidal and retinal degenerations without CNV, including geographic atrophy in dry AMD, or to test cell replacement of choroidal endothelial cells toward potential human therapies in a wide range of disease states.

## Acknowledgements

We would like to thank the University of Iowa Animal Care staff for their help with the project and their commitment to ensuring the fair and ethical utilization of animals for important biomedical research.

## Supplementary Information

**Supplementary Figure 1.**
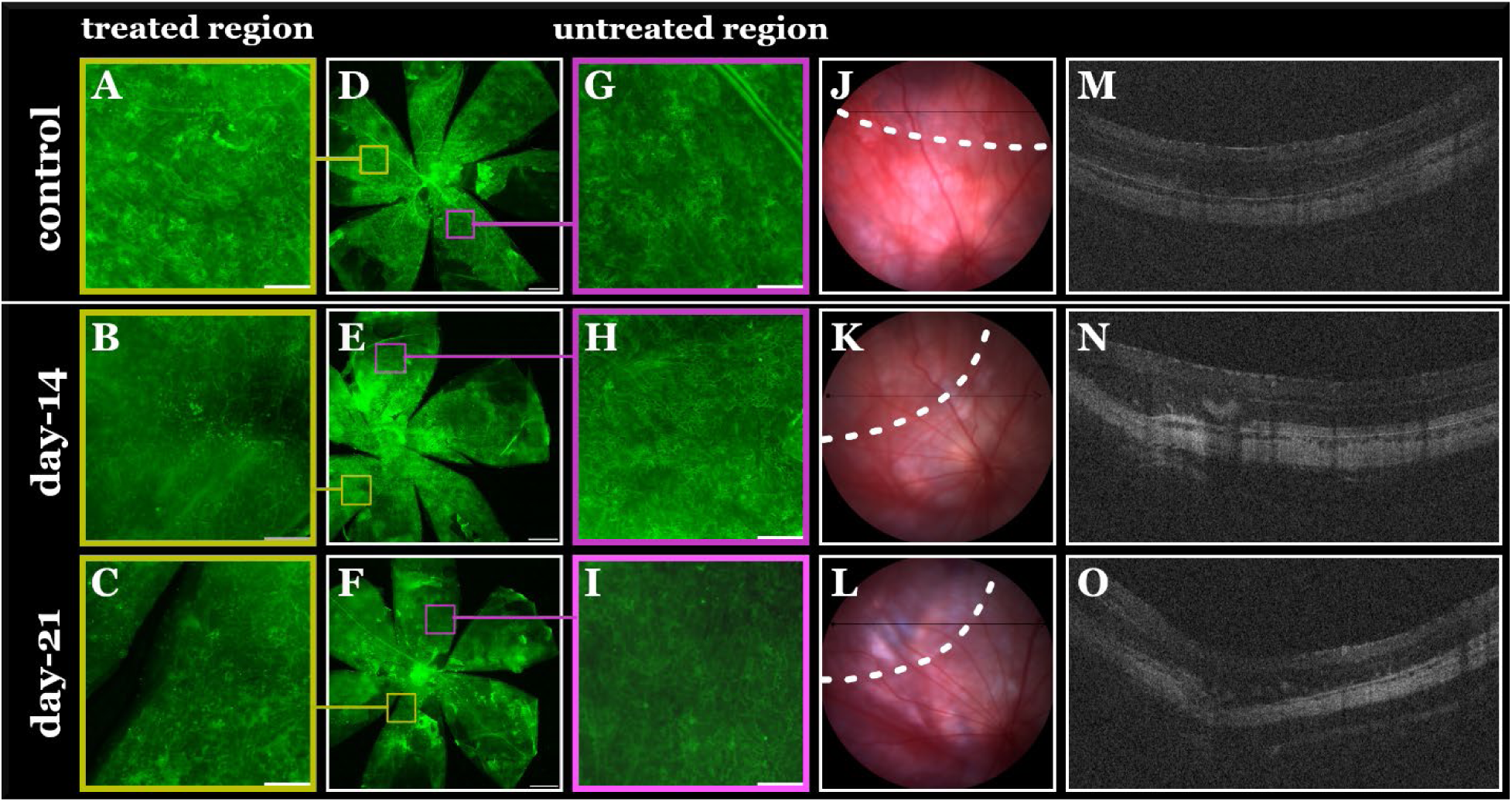
Suprachoroidal anti-CD105-SAP injection causes sectoral choroidal and outer retinal loss. Suprachoroidal injection of anti-CD105-SAP (0.5 µg/eye) causes sectoral choroidal and outer retinal loss compared to eyes injected with control (PBS only; top row). High magnification inserts (1st and 3rd column; scale bar 100 µm) from flat mounts (D-F; Scale bar is 1000 µm) demonstrate focal choroidal injury in treated areas (A-C, Scale bar is 100 µm) and preserved choroidal vasculature in untreated areas (G-I, Scale bar is 100 µm). Corresponding fundus photographs, with the site of injection, showing in white dashed lines(J-L) and OCT line scans (M-O) show sectoral regions of choroidal and outer retinal damage, which persists through all studied time points.

**Supplementary Figure 2.**
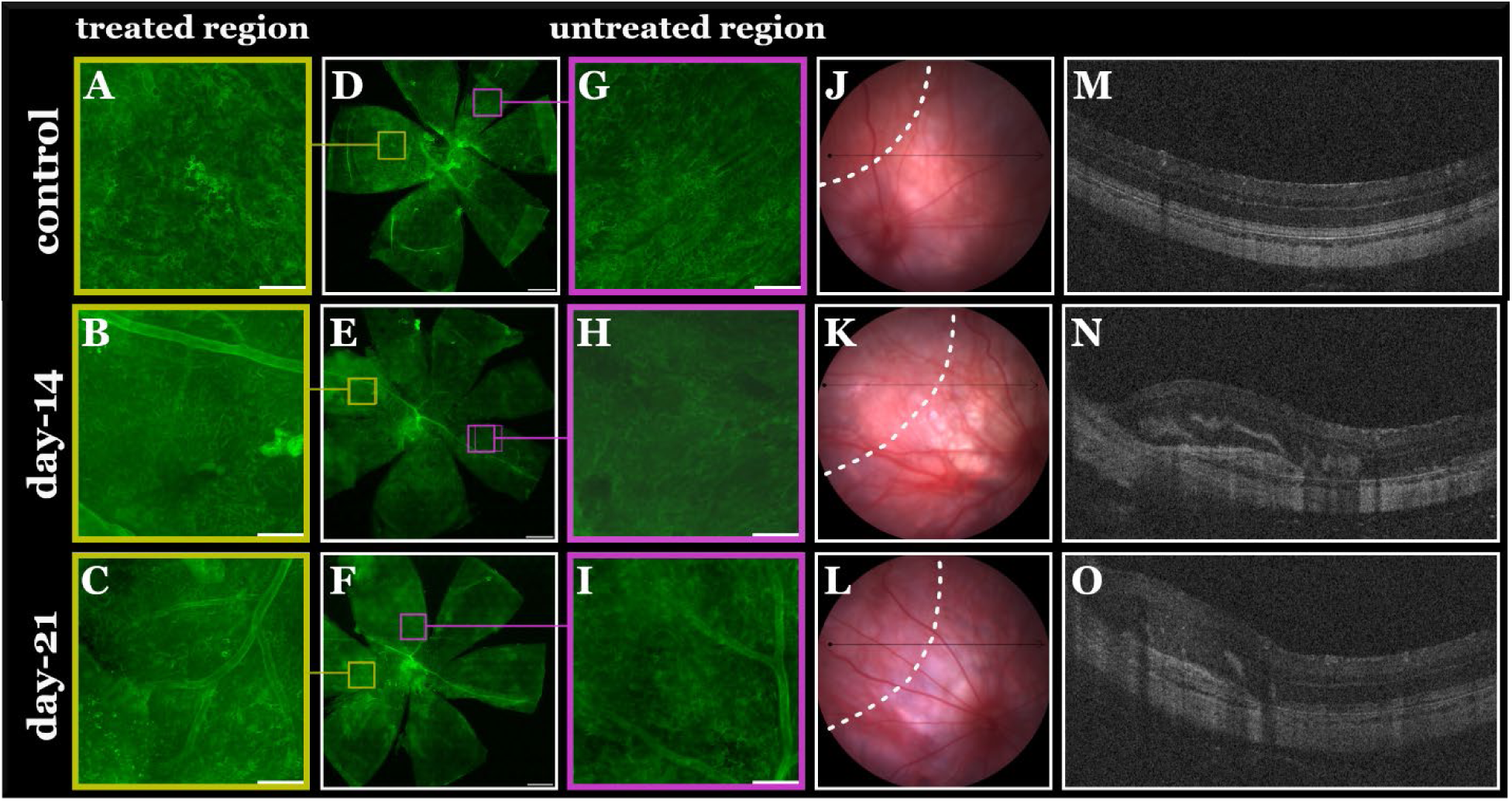
Suprachoroidal anti-CD38-SAP injection causes sectoral choroidal and outer retinal loss. CD38-SAP suprachoroidal injection leads to sectoral loss of choroidal and outer retinal tissue compared to eyes that received the control injection (PBS only; top row). High magnification images (1st and 3rd column; scale bar 100 µm) from choroid scleral flat mounts reveal localized choroidal damage in the treated regions (A-C, Scale bar is 100 µm), while the choroidal vasculature remains intact in untreated areas (G-I, scale bar is 100 µm). Corresponding fundus images, with the site of injection, showing in white dashed lines (J-L) and OCT scans (M-O) display distinct sectors with choroidal and outer retinal damage.

**Supplementary Figure 3.**
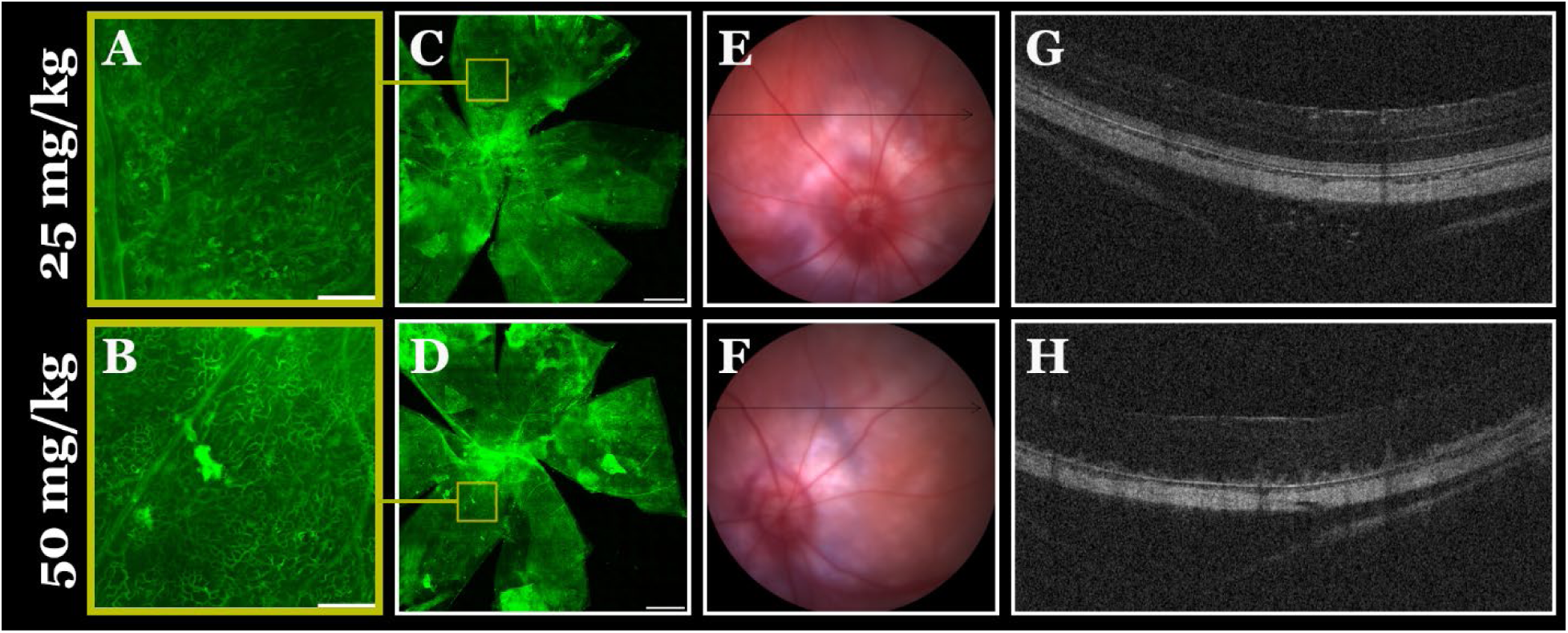
Systemic administration of sodium iodate in SRG rats results in dose-related choroidal and outer retinal damage at 1-week post-treatment. Intravenous sodium iodate resulted in diffuse choroidal and outer retinal loss in a dose-related fashion (doses shown on left) by 1 week after treatment. High magnification inserts (A,B; Scale bar is 100 µm) from flat mounts (C,D); Scale bar is 1000 µm) demonstrate choroidal injury. Corresponding fundus photographs (E,F) and OCT line scans (G,H) show diffuse choroidal and outer retinal damage. Note the minimal choroidal and outer retinal damage in low-dose treated animals (25 mg/kg; top row).

